# Absence of electroencephalographic evidence of implicit action-effect intentional binding with varying effect probability

**DOI:** 10.1101/2020.12.17.423230

**Authors:** Max Seignette, Mark Schram Christensen

## Abstract

The subjective experience of an attraction in time of an action, and the event caused by the action, is known as the intentional binding phenomenon. Intentional binding is a robust phenomenon and has previously been associated with subjective sense of agency, but recent studies have shown that binding can take place in the absence of action intentions. In this study, we tested possible electrophysiological equivalents to the intentional binding phenomenon under a simple action-effect task, where pressing of a button caused tones to occur at different pitches or delays with different probabilities. Changing the probabilities of the effect of an action has in some previous studies shown to influence the intentional binding phenomenon. We tested whether changes in action-effect probability gave rise to differences in movement related cortical potentials (MRCP) slopes, peak latency and auditory event related potential (aERP) changes of amplitude or latency of the N1, P2, P3 and N4 components of the central aERP, of which some has been related to sense of agency or intentional binding. We also tested differences in MRCP across the whole scalp prior to movements, and to differences in aERP across the whole scalp after the tone is played. We found no electrophysiological indications of intentional binding when action-effect contingencies were changed in accordance with conditions that have given rise to intentional binding in previous experiments. Our results are in line with several recent studies that have questioned whether intentional binding follows all voluntary actions and can be related to sense of agency at all.

## Introduction

The intentional binding phenomenon is a robust, reproducible finding, which shows a perceptual attraction in time of a voluntary action and the subsequent event caused by the action. According to the intentional binding principle (Haggard et al. 2002), the subjective judgment of when an action is performed and when the effect is experienced are attracted to each other in time, when one is in a situation where one performs the action oneself. This attraction in time of the experience of the action and experience of the consequence has been related to the sense of agency and suggested to function as an implicit measure of the sense of agency, which can be studied independent of making explicit judgments of one’s subjective experiences of control of own movements. It has previously been argued that human voluntary actions (always) are followed by a sense of agency, such as exemplified in quotes like: “we experience a clear feeling (or ‘buzz’) of agency during everyday actions, even when no evaluation or judgement is required” (Haggard (2017), p. 199), and that this sense of agency seems to be at the core of the intentional binding phenomenon, as exemplified by Blakemore and Frith (2003, p. 220): “The temporal attraction between self-generated actions and their sensory consequences binds together these events, and enhances our experience of agency’’. So regardless of whether it is measured or not, the feeling of agency and its accompanying phenomenon of intentional binding should always be present according to these arguments. This has been the point of departure for the present experiments.

Nonetheless, we also acknowledge that the concept of a sense of agency, and its accompanying intentional binding phenomenon, have not been undisputed. Several previous studies many of them recent, such as Saito et al. (2015), Schwarz et al. (2019), Kirsch et al. (2019) and Ma et al. (2020) have explicitly investigated the relationship between the intentional binding phenomenon and the subjective experience of sense of agency and found no correlation.

However, Imaizumi & Tanno (2019) showed that explicit measures of agency and intentional binding were correlated. The method used by Imaizumi & Tanno (2019) with increasing delays between action and effect to diminish sense of agency and intentional binding, has previously shown to increase intentional binding (Wen et al. 2015). This suggests that the method of increasing delays alone to decrease sense of agency is not a straightforward method and that context and prior experience with the exact action-effect relations may be of importance for the interpretation of this method. Dewey & Knoblich (2014) showed that two different indirect measures of agency, intentional binding and sensory attenuation, were not correlated. It has also been suggested that the intentional binding phenomenon reflects causal beliefs, for instance that lightning causes thunder, rather than subjective experience of agency (Desantis et al. 2011; Ma et al. 2020). Based on these studies it is not possible to determine whether intentional binding and sense of agency measure the same underlying phenomenon. But the number of studies questioning the relationship between intentional binding and sense of agency may suggest that the two mechanisms are independent of one another.

Regardless of whether intentional binding reflects sense of agency or the causal belief, that an action leads to an effect, the intentional binding should also be present in the absence of an explicit enquiry into the intentional binding measurement. Since the intentional binding phenomenon is a subjective attraction of an action and the following event, it is expected that a neural correlate to the intentional binding phenomenon should be present irrespective of the actual measurement of the intentional binding – under the assumption that a subjective experience can be reflected as a neural correlate.

In this study, we would like to investigate indirectly the argument presented in the beginning, that our actions are accompanied with a feeling of agency. We will investigate this assumption using an action-effect manipulation paradigm that previously has been used to influence the intentional binding phenomenon. This was accomplished by manipulating with the probability of the outcomes of actions, even when the action is produced by oneself in all situations. Engbert & Wohlschläger (2007) showed that changes in the probability of whether a tone was played or not played after a button press had an impact on the intentional binding phenomenon. The intentional binding phenomenon was only present when there was a higher probability of the action causing a tone in 2/3 of trials. When there was a lower probability, i.e. the action only caused a tone in 1/3 of the trials, the binding phenomenon was abolished. In a study by Moore and Haggard (2008), a tone occurred with either 50% or 75% probability after a button press, which gave rise to a shift in the perception of the occurrence of the button press, in sessions where the tone occurred with a 75% probability. In a similar study, Moore et al. (2009) asked participants to evaluate if an action occurred when it was followed by a tone in 75% or 50% of cases, and again found a shift in the perception of the action towards the tone. The same procedure was also used by Voss et al (2010) with similar findings.

These studies suggest that by manipulating statistical probabilities of outcomes of specific actions, one can influence the intentional binding phenomenon. Consequently, we may also be able to tap into the neural mechanisms underlying the intentional binding phenomenon using measurements of cortical electrical activity from electroencephalography (EEG) recordings. If the intentional binding reflects a real perceptual or psychological phenomenon, one expects that neural mechanisms also would reflect the binding phenomenon. These could include attraction of specific neural signatures in time, changes in peak amplitude or latency of specific EEG components related to action and effect or differences in coupling patterns between brain areas.

In terms of electrophysiological markers related to the intentional binding phenomenon, a couple of candidates have been suggested. Jo et al. (2014) showed that the slope of the early part of the readiness potential (RP) (Kornhuber & Deecke, 1965; Shibasaki & Hallett 2006) reflect differences in the intentional binding phenomenon. In particular, the slope of the early part of the RP showed a positive correlation with perceived shifts of the time of the tone. This was reflected in such way, that the large negative slopes of the early RP were associated with an earlier temporal perception of the tones.

We assume that EEG signals reflect underlying cognitive or psychological states associated with one’s subjective reaction to uncertainty. Therefore, it is reasonable to suggest that EEG signals may reflect the underlying statistical contingencies of action-effect outcomes when these are changed in systematic ways. In the present study we therefore manipulated with action-effect outcomes such that the outcome either appear with a low or high degree of uncertainty. Both Engbert & Wohlschläger (2007), Moore and Haggard (2008) and Moore et al. (2009) have shown that manipulations of outcome uncertainty influence intentional binding. If we combine that knowledge with the results from Jo et al. (2014), i.e. that the slope of the early RP correlates with action binding effects, we expect that our manipulation of the action-effect probability gives rise to different neural signatures in situations with a low or high degree of uncertainty.

If high certainty action-effects contingencies are more prone to give rise to intentional binding compared with low certainty contingencies, we would expect that the slope of the early RP is different in those two situations. In this study we would therefore like to test the idea, that if one has been exposed to high certainty action-effect contingencies through several trials, one has learned what the consequences of one’s actions are, which will be reflected as neural marker in the EEG, in particular the slope of the RP. Vercillo et al. (2018) have also shown that changing the action-effect contingency in terms of different sensory modalities, where an action causes a visual effect or a visuo-motor effect, have an impact on the amplitude of central readiness potential, providing different amplitudes of the Cz electrode voltage prior to the action.

Another line of electrophysiological evidence that is relevant in relation to action-effect outcome probability, is the modulation of auditory event related potentials (aERPs) depending on the predictability of the outcome tones. Schafer & Marcus (1973) showed that the amplitude of aERPs is lower when one generates a tone oneself, compared with externally generated tones. Furthermore, they also showed that predictable tones show lower aERP amplitudes than unpredictable tones. A study by Timm et al. (2016) showed that the both the N1 and P2 components of aERPs were reduced in amplitude when tones were self-generated, and that the amplitude of the P2 correlated with subjective sense of agency. This suggest that the P2 component of aERP is of special interest to study electrophysiological markers of sense of agency, which the intentional binding phenomenon has been suggested to reflect. We would therefore expect that situations where an action leads to a predictable effect with high probability, the P2 component would be reduced in amplitude compared to a situation where the action-effect outcome is less predictable. Finally, the P3 component of aERPs have also been investigated in relation to action-effect studies (Vastano et al 2020) and stimulus response studies (Kleimaker et al. 2020; Takacs et al. 2020; Takacs et al. 2021). Therefore, we will also investigate whether the predictability of the outcome tone changes the P3 component.

A third line of electrophysiological evidence indirectly related to the intentional binding phenomenon is the use of dynamic causal modelling (DCM) for induced responses (Chen et al. 2008) on a study of sense of agency. Ritterband-Rosenbaum et al. (2014) showed that in the late phase of visuomotor line drawing movements, when participants reported a positive sense of agency of the movement, oscillations in the 50-60 Hz range in the inferior parietal cortex modulated oscillations in the 40-70 Hz range in preSMA. If intentional binding and sense of agency share a common neural mechanism, one would expect that a situation that leads to intentional binding would share the same coupling patterns between IPC and preSMA as found by Ritterband-Rosenbaum et al. (2014). On the other hand, studies which have shown the intentional binding phenomenon can all be characterised as action-effect studies (see Christensen & Grünbaum 2017; Christensen & Grünbaum 2018), whereas the study by Ritterband-Rosenbaum et al. (2014) is an example of a feedback manipulation study. Therefore, there may be some differences in neural correlates depending on whether the study is an action-effect or a feedback manipulation study. However, that may indicate that action-effect studies and feedback manipulations do not share underlying mechanisms of sense of agency. A positive sense of agency, regardless of whether it is studied as a feedback manipulation experiment or action-effect experiment, has among other regions been associated with increased activation measured with functional magnetic resonance imaging in precuneus (David et al. 2007, Spaniel et al. 2016, Fukushima et al. 2013) and supplementary motor area (David et al. 2007, Farrer et al. 2008, Kühn et al. 2013). In contrast, an explicitly reported lack of sense of agency or perception of incongruency between movement and feedback has been associated with changes of activity in left dorsolateral prefrontal cortex (Farrer et al. 2008, Spaniel et al. 2016, Chambon et al. 2013, David et al. 2007, Kontaris et al. 2009, Nahab et al. 2011) and left angular gyrus (Farrer et al. 2002, Farrer et al. 2008, Chambon et al. 2013). However, testing multiple potential network models derived from different fMRI experiments using DCM for induced responses in EEG is beyond the scope of this study. Therefore, we have focused on EEG findings that are related to potential differences between conditions, which should or should not give rise to either sense of agency or intentional binding.

To test the existence of electrophysiological correlates to the intentional binding phenomenon independent of measurements of intentional binding, we therefore adopted the idea of manipulating action-effect outcome probabilities similar to at least two different previous experimental setups (Engbert & Wohlschläger 2007; Moore and Haggard (2008); Moore et al. (2009); Voss et al. (2010)). These studies have shown that manipulating action-effect outcome probabilities influence the intentional binding effect. Therefore, we investigated situations where the consequences of a button press lead to different tone characteristics. In our experiment we manipulated the occurrence probability of two tones in two different experimental sessions, in a 50/50 % distribution and in a 20/80 % distribution. In one experiment, tones that followed a button press with a constant delay of 250 ms could occur at two different pitches (high or low pitch). In the other experiment, the same low pitch tone could occur after delays of either 250 ms or 600 ms after the button press. Each of the two experiments was conducted in random order and within each experiment the conditions (20/80 or 50/50) were performed in random order.

At the same time, we recorded EEGs from the participants to investigate whether it was possible to explore and detect any differences in the EEG measures from the time prior to the action and until after the outcome tone was played.

To investigate action-effect contingencies in the described experimental setup, we tested the following hypotheses: Concerning RPs, we expected that the slope of the early RP would be different in the 20/80 conditions compared to 50/50 conditions. Furthermore, we predict that the aERPs, in particular the P2 following the tones, for two identical tones would be shifted towards the time of the button press in the high certainty 20/80 conditions compared to the uncertain 50/50 condition. Besides these specific differences in EEG measurements, we further explore differences between the 20/80 and 50/50 conditions using scalp-time maps both prior to and during the button presses and following the tones. Finally, we also performed Dynamic Causal Modelling (DCM) analyses for induced responses of the coupling between IPC and preSMA between the button press and the tone comparing the 20/80 and 50/50 conditions.

## Methods

### Participants

Twenty-three healthy participants, of which seven were female, were recruited for this study. Participants ranged in age from 18 years to 50 years, mean age was 24 years, standard deviation 4.8 years. Participants were recruited through an online participant-recruitment site: www.forsogsperson.dk and through advertisements at the Faculty of Health Sciences at University of Copenhagen. All participants received oral and written information about the study and were given 24 hours to decide whether they would provide their written informed consent, which all of the participants did. The study was approved by the local ethics committee of the Capital Region of Copenhagen, Region H (protocol number: H-3-2013-198).

### Experimental procedures

Participants were placed in a dark room, where the only light source was the two computer screens used to control the recording equipment. Participants were seated in a chair 1.5 m from a wall, which they were facing, and they were instructed to look at a fixation cross on the wall. Each participant underwent two experiments (Delay and Pitch experiments) that in total comprised four experimental sessions, all taking place sequentially on the same day, in semi-random order. In all four sessions, participants were instructed to press a button with their right thumb at their own pace for 150 times (but not faster than one button press every 3 s). The button press gave rise to an auditory tone played from a loudspeaker placed ∼1,5 m behind the participant. Across all participants, the shortest session lasted 566 s and the longest 2249 s calculated from first button press to last tone.

### Pitch experiment

In two of the sessions, the tones presented were either a low pitch tone (220 Hz) or a high pitch (440 Hz) tone, each lasting 100 ms. The tones were presented 250 ms after the participant had pressed the button. In one of the conditions the high and low pitch tones were presented in random order with an equal probability, i.e. 50% high pitch, 50% low pitch. In the other of the two sessions, the low pitch tone was presented in 80% of the trials and the high pitch in 20% of the trials in random order.

### Delay experiment

In the other two sessions a low pitch tone (220 Hz) was presented exclusively, but it was either presented with a delay of 250 ms or 600 ms after the button press. In one of these sessions the probability of the delays was distributed equally (50%/50%) between the two, in the other session the 600 ms delay was presented in 20% of the trials and the 250 ms delay in 80% of the trials in random order.

The order of the four sessions was controlled in such a way that both sessions in one experiment was performed after each other, but in pseudo random order, and the order of the Delay experiment or Pitch experiment was also pseudo random. I.e., either the two pitch sessions were presented first, or the two delay sessions were presented first in random order.

The pressed button was connected to a 9V battery and gave rise to a square voltage pulse, which was detected by a Cambridge Electronic Design Micro 1401 Mark2 AD converter (Cambridge Electronic Design Ltd., Cambridge, United Kingdom). The detection of the up-going flank of the TTL pulse triggered a frame in the software program Signal 5.2 (Cambridge Electronic Design Ltd., Cambridge, United Kingdom) which provided the signal that generated the tones played through loudspeakers placed behind the participant.

The objective with the different probability conditions was to create situations in which the 20%/80% probability distribution, the action (pressing the button) lead to an effect that was presented with high certainty (80%), and the tone with low probability became a surprising event. In the 50%/50% condition, the effect of the action became highly uncertain, because the probability of each effect was equal. By the last 2/3 part of the different sessions, we assume that the participants have become acquainted with the situation of being in a high probability certain situation (in the 20/80 situation) or in a low probability uncertain situation (50/50 situation).

### Electroencephalogram (EEG) recordings

Before participants performed the four experimental sessions, they were equipped with a 64 electrode EEG cap attached with BioSemi Active2 active electrodes and amplifier (BioSemi, Amsterdam, The Netherlands). EEG was recorded using the software ActiView6.5 (BioSemi, Amsterdam, The Netherlands) EEG was recorded at 2048 Hz in bdf format files. Triggers indicating button presses and auditory tones were provided to the data file using digital output signals from the Micro 1401 AD converter. The EEG data were stored on a laptop computer for subsequent analyses. Data acquisition from one participant was stopped, due to insufficient data quality, which was not possible to adjust (using ICA and filtering). Data from that participant (Participant 11) was omitted from data analysis.

### Behavioural analyses

For all four conditions, the interval between each of the button presses (inter press interval, IPI) were calculated based on the information about when the button press triggers were detected in the EEG data. Subsequently, we compared the IPI in the four conditions using a within-subject analysis of variance with the two factors: Experiment (Delay vs Pitch) and Probability Condition (higher 20/80 vs lower 50/50) (See Figure 1A, 1D). In addition, we also sorted button presses according to whether they followed long or short delay tone in the Delay experiment or followed high or low pitch tones (Figure 1B,1C) in the Pitch experiment (Figure 1E,1F).

**Figure 1:**
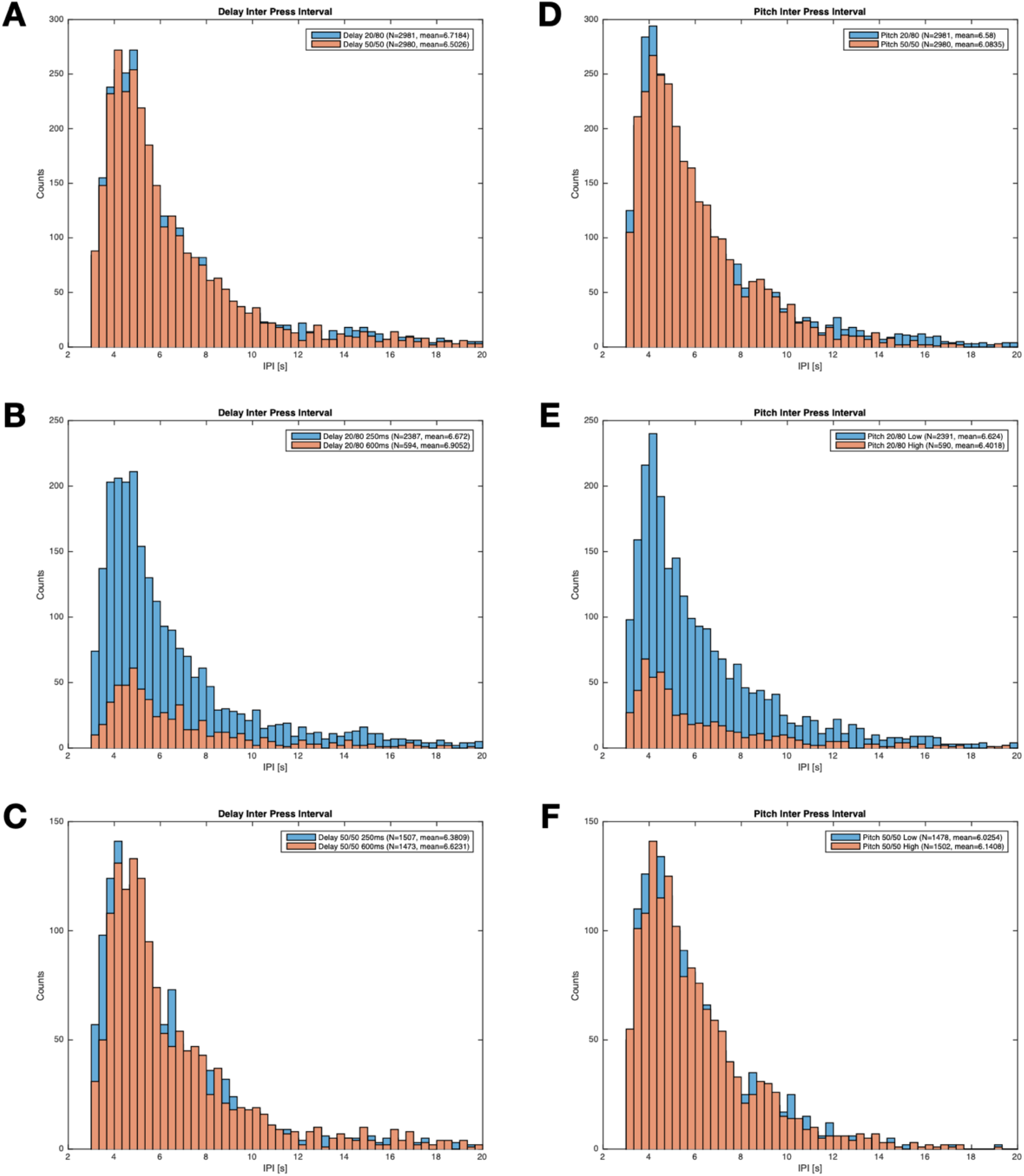
Inter press intervals (IPI). **A** shows two histograms for all trials of the IPI for the Press Delay condition. In blue the 20/80 probability condition, in orange the 50/50 probability condition. **B** shows the histogram of the IPI in the Delay 20/80 condition, where IPIs are sorted according to whether they follow a high probability 250ms delay tone or a low probability 600 ms delay tone. **C** shows the histogram of the IPI in the Delay 50/50 condition sorted into whether they follow long or short delays. **D** shows histograms for all trials for the Press Pitch condition. In blue the 20/80 probability condition, in orange the 50/50 probability condition. **E** shows the histogram of the IPI in the Pitch 20/80 condition, where IPIs are sorted according to whether they follow a high probability low pitch tone or a low probability high pitch tone. **F** shows the histogram of the IPI in the Pitch 50/50 condition, where IPIs are sorted according to whether they follow a low pitch tone or a high pitch tone.

### EEG analysis

EEG files from the four different sessions were initially imported into the software EEGlab (v13.4.4b) (https://sccn.ucsd.edu/eeglab/, Swartz Center for Computational Neuroscience, La Jolla, CA, United States) for Matlab (R2015a) (MathWorks, Natick, MA, United States), where default channel locations for the BioSemi electrode cap was applied. For computational optimisation reasons, the data were down-sampled from 2048 Hz to 256 Hz. For analyses of movement related cortical potentials (MRCP), in particular lateralized readiness potentials (LRPs), central readiness potentials (RPs) and -10 ms movement related potentials (N-10), see Shibasaki & Hallett (2006), a high pass filter of 0.05 Hz was applied. For analysis of aERPs, a separate copy of the data was made where a high pass filter of 1 Hz was applied. These filters were applied to the down-sampled data. The subsequent pre-processing steps of the MRCPs and aERPs were similar, only when we moved to more advanced analyses, the two differed again. The MRCP and aERP data were then low pass filtered with an 80 Hz filter and the data were re-referenced to an average reference, excluding EEG channels with obvious noise contamination, typically 50 Hz line noise. An independent component analysis (ICA) was then employed using the runICA algorithm implemented in EEGlab. ICAs revealing eye-blinks, lateral eye-movements, and muscle activity were excluded from the data.

Data were then imported into SPM12 (http://www.fil.ion.ucl.ac.uk/spm/, Wellcome Trust Centre for Neuroimaging, University College London, London, United Kingdom), where they were epoched based on the time point of the button press indicated by a trigger for MRCP analyses from -2500 ms before the time of press until +1000 ms after the button press. Data for the MRCP analyses were baseline corrected using the average of the individual channel potentials calculated from -2500 to -2000 ms before the button press. Alternatively, epoching was based on the time point of the presentation of the auditory tone indicated by another trigger for the aERP analyses, again from -1000 ms to +1000 ms around the time of the auditory tone. In order not to mix effects of the MRCP into the baseline correction, the aERP data were baseline corrected using the average of the individual channel potentials calculated from 750 to 1000 ms after the auditory tone was presented.

All press epochs from the 50/50 and 20/80 Delay experiment conditions were combined into a single file, giving rise to two different Press Delay conditions: Press Delay 50/50 and Press Delay 20/80.

All press epochs from the 50/50 and 20/80 Pitch experiment conditions were combined into a single file, giving rise to two different Press Pitch conditions: Press Pitch 50/50, Press Pitch 20/80.

All tone epochs from 50/50 and 20/80 Delay sessions were combined into a single file, giving rise to four different Tone Delay conditions: Tone Delay 250ms 50/50, Tone Delay 600ms 50/50, Tone Delay 250ms 20/80, Tone Delay 600ms 20/80.

All tone epochs from 50/50 and 20/80 pitch sessions were combined into a single file, giving rise to four different Tone Pitch conditions: Tone Pitch 50/50 low, Tone Pitch 50/50 high, Tone Pitch 20/80 low, Tone Pitch 20/80 High.

To investigate neuronal responses in the situations where the participants have become acquainted with the higher probability (20/80) and lower probability (50/50) consequence of the action, we used the first 50 trials in each session to serve as adaption trials to the statistical contingencies employed in the given round. In a two-action two-effect experiment Desantis and Haggard (2016) used 40 or 20 trials for learning action-effect associations, so 50 trials should serve as more than sufficient to learn that pressing a button most like lead to a specific tone in the high 20/80 probability conditions. Subsequently, we only performed analyses on the last 100 trials in the different sessions, similar to what has been used in previous analyses comparing the effect of action-effect contingencies on RPs and visual evoked potentials in a different task (Vercillo et al. 2018). This gave a reasonable balance between having learnt the relations between actions and events, and at the same time enough trials to analyse for MRCPs and aERPs.

Averages for the four different (Press Delay 20/80, Press Delay 50/50, Press Pitch 20/80 and Press Pitch 50/50) LRPs and RPs were calculated based on the last 100 trials and C3 electrode responses were calculated for each participant. We did not employ formal statistical tests of neither onset nor amplitude of the LRPs/RPs. We calculated the slope of the early and late LRP/RP as described in Jo et al. (2014). These were calculated from an average signal of the 9 electrodes around Cz. Briefly, the slope of the early LRP/RP is calculated as the slope from -2500 to -1000 ms and the late is between -500 ms and 0 ms. The signal values used to calculate the slope at the different point are determined as the difference of the average signal amplitude from (2500-2300 ms before button-press) compared with the average signal amplitude from (1000-800 ms before button-press), from (700-500 ms before button-press) and from (200-0 ms before button-press). In Figure 2 the early and late slopes of the LRPs and ERP have been sketched in the figures. Furthermore, we calculated the peak time and amplitude of the late phase of both the LRP and RP as the time point and amplitude of the maximal negative potential between -100 ms and 100ms around the time of the button press, which by definition is set to 0 ms, also known as the N-10 component. These responses are displayed in Figure 2.

**Figure 2:**
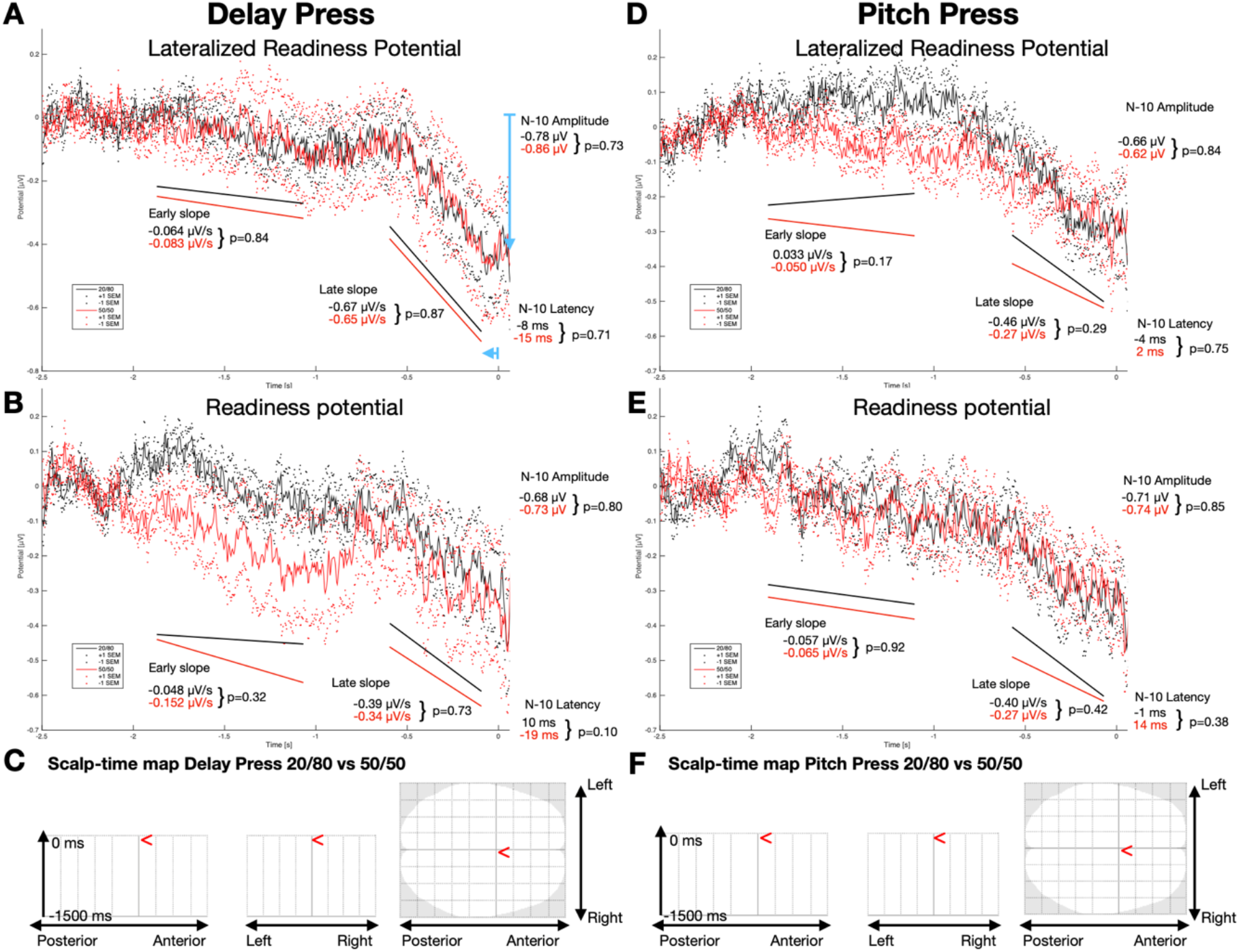
Readiness potentials of the different conditions in the two experiments. **A** LRP from an average of 9 electrodes centered at C3 for the Delay Press condition. Early and late slopes were calculated as the means of averages of the individual LRP, and may diverge form the actual slope of the grand mean averages displayed in the graphs. Therefore, slopes are not present overlying with graphs but underneath. **B** central readiness RP from an average of 9 electrodes centered at Cz for the Delay Press condition. **C** Scalp-time map of the difference in scalp potential of the 20/80 and 50/50 conditions of the Delay press experiment from -1500 ms to 0 ms from button press. SPM{F} map is initially thresholded at p<0.001 uncorrected and then clusters of activated pixels that survive a correction for multiple comparisons p<0.05 FWE are depicted. No areas were significant. **D** Lateralized readiness potential from an average of 9 electrodes centered at C3 for the Pitch Press condition. **E** central readiness potential from an average of 9 electrodes centered at Cz for the Pitch Press condition. F Scalp-time map of the difference in scalp potential of the 20/80 and 50/50 conditions of the Pitch press experiment from -1500 ms to 0 ms from button press. SPM{F} map is initially thresholded at p<0.001 uncorrected and then clusters of activated pixels that survive a correction for multiple comparisons p<0.05 FWE are depicted. No areas at any time were significant.

Averages for the eight different (Tone Delay 250 ms 20/80, Tone Delay 600 ms, 20/80 Tone Delay 250 ms 50/50, Tone Delay 600 ms 50/50, Tone Pitch 20/80 Low, Tone Pitch 80-20 High, Tone Pitch 50-50 Low, Tone Pitch 50-50 High) aERPs were calculated based on the last 100 trials and Cz electrode responses for each participant. For the 20/80 probability conditions that gave 20 trials to calculate ERPs from the low probability tones and 80 trials to calculate ERPs from the high probability. In the 50/50 probability condition, 50 trials were used to calculate the average ERPs. We performed statistical tests of the difference of the latency and amplitude of the N1, P2, P3 and N4 aERP component between the 20/80 and 50/50 conditions using paired t-tests for the 250 ms delay tones in the Delay Tone experiment, and of the low pitch tones in the Pitch Tone experiment. The N1 ERP component was found as the minimal negative deflection of the average signal from the 9 electrodes around and including Cz in the time interval from 70 ms to 140 ms, the P2 was found as the maximal positive deflection of the average signal from the 9 electrodes around and including Cz in the time interval from 135 ms to 265 ms, the P3 was found as the maximal positive deflection of the average signal from the 9 electrodes around and including Cz in the time interval from 250 ms to 350 ms, and the N4 ERP component was found as the minimal negative deflection of the average signal from the 9 electrodes around and including Cz in the time interval from 350 ms to 500 ms,

Data from one of the remaining participants were discarded for group analyses because no clear MRCPs or aERPs were identifiable on averages from this participant (Participant 20). Finally, one participant (Participant 6) was excluded from further analyses due to excessive frontal electrode amplitudes despite high pass filtering and exclusion of ICA components related to eye-blinks.

For subsequent group analyses the total number of participants was twenty (n=20).

Grand mean averages were calculated for the four different MRCPs (Figure 2A and 2C) and eight different aERPs (Figure 2D and 2E).

Finally, we also calculated an electrophysiological equivalent to the intentional binding by calculating the difference in the latency of the negative part of the MRCP around the time of the button press (N-10 component) and N1 and P2 components of the aERPs.

### Scalp-time EEG analysis

To formally tests differences in MRCPs at any location on the scalp between the Press Delay 20/80 and Press Delay 50/50 conditions (Figure 2C) and the Press Pitch 20/80 and 50/50 conditions (Figure 2) the 64 channel EEG averages from each participant was filtered with a 30 Hz low pass filter and interpolated into scalp-time (i.e. 2D-time) maps. The maps are 2D scalp images of the EEG amplitudes with a temporal dimension, meaning that at each time sample of the 64 channels a 2D image is interpolated into a 32×32 grid. The 2D scalp images are a projection of the 3D locations of the 64 channels onto a 2D slightly skewed ellipse and the image intensity is the EEG amplitude in the 64 channel locations and interpolated in the loci between the actual channel locations. These maps were then constructed in a limited time window between -1500 ms and 0 ms adjusted to the button press. These scalp-time maps entered a 2nd level group analysis in the form of a repeated measures one-way ANOVA, with the factor 20/80 versus 50/50 conditions.

To formally test the difference between the four different tone conditions across the whole scalp in the two different experiments, the 64 channel EEG averages from each participant was filtered with a 30 Hz low pass filter and interpolated into scalp-time (i.e. 2D-time) maps. These scalp-time maps were constructed of the time interval from the time of the appearance of the tone until 600 ms after the tones were played.

For the Tone Delay (Figure 4 A-D) and Tone Pitch (Figure 4 E-H) experiments the scalp-time maps entered a 2nd level group analysis in the form of a repeated measure two-way ANOVA with the factors 20/80 vs 50/50 and High vs. Low Pitch or 250 ms/600ms Delay.

**Figure 3:**
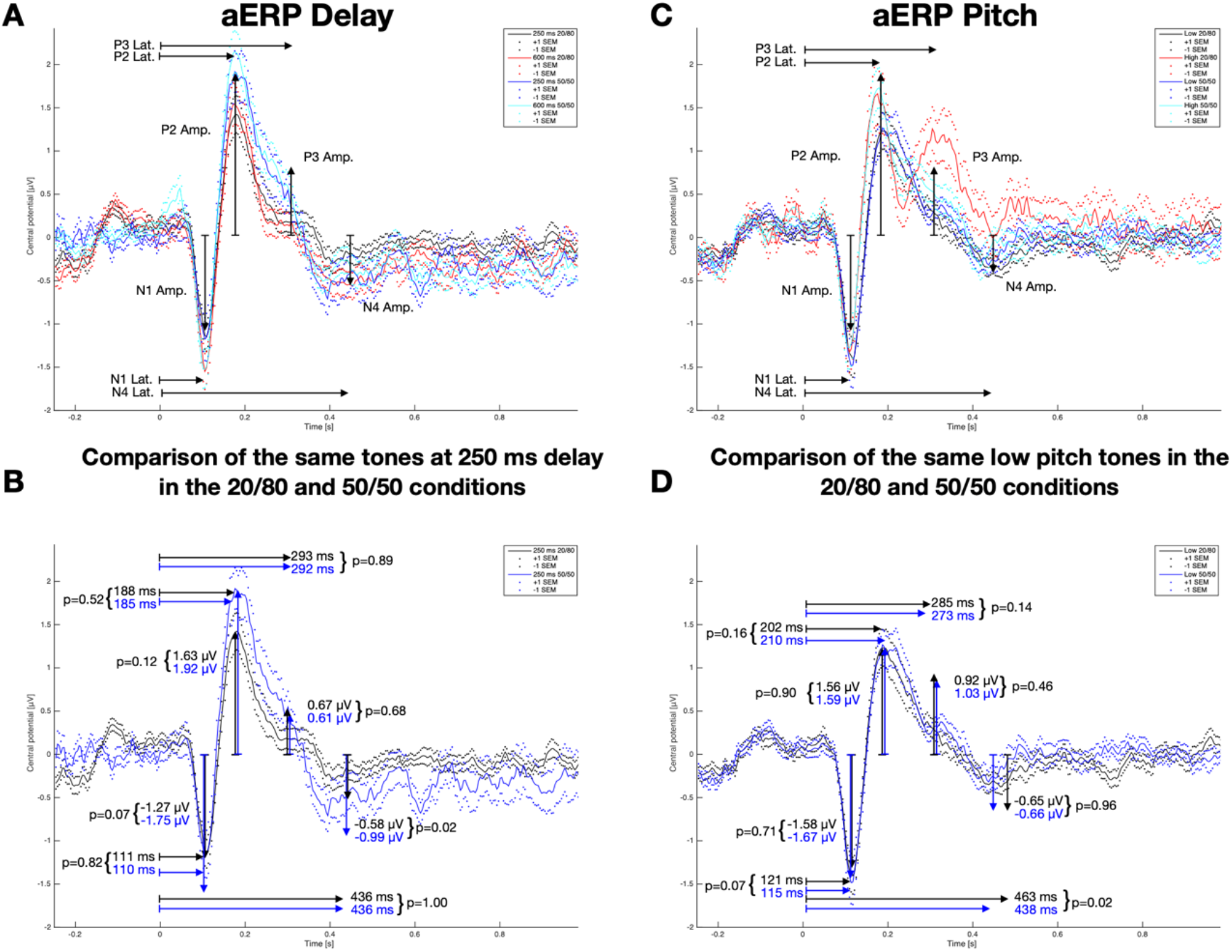
Auditory event related potentials of the different tones in the two experiments. **A** shows the aERP of the Delay Tones, in black: 250 ms delay tones in the 20/80 condition, red: 600 ms delay tones in the 20/80 condition, blue: 250 ms delay tones in the 50/50 condition, cyan: 250 ms delay tones in the 50/50 condition. **B** shows the two comparable tones both at 250 ms delay after the button press in either the 20/80 or 50/50 condition. Values of N1, P2, P3 and N4 latency and amplitude are displayed for both conditions. These are calculated as the mean of the N1, P2, P3 and N4latencies and amplitudes derived from the individual participant’s N1, P2, P3 and N4 latency and amplitude. The graphs of the aERPs are calculated as grand mean averages of the aERPs of all participants, and may therefore differ a little from the displayed numbers. P-values of paired t-tests are displayed along with the values. **C** shows the aERP of the Pitch Tones, in black: low pitch tones in the 20/80 condition, red: high pitch tones in the 20/80 condition, blue: low pitch tones in the 50/50 condition, cyan: high pitch tones in the 50/50 condition. **D** shows the two comparable low pitch tones after the button press in the 20/80 and 50/50 conditions with displayed latency and amplitude values and p-values of the paired t-test comparisons.

**Figure 4:**
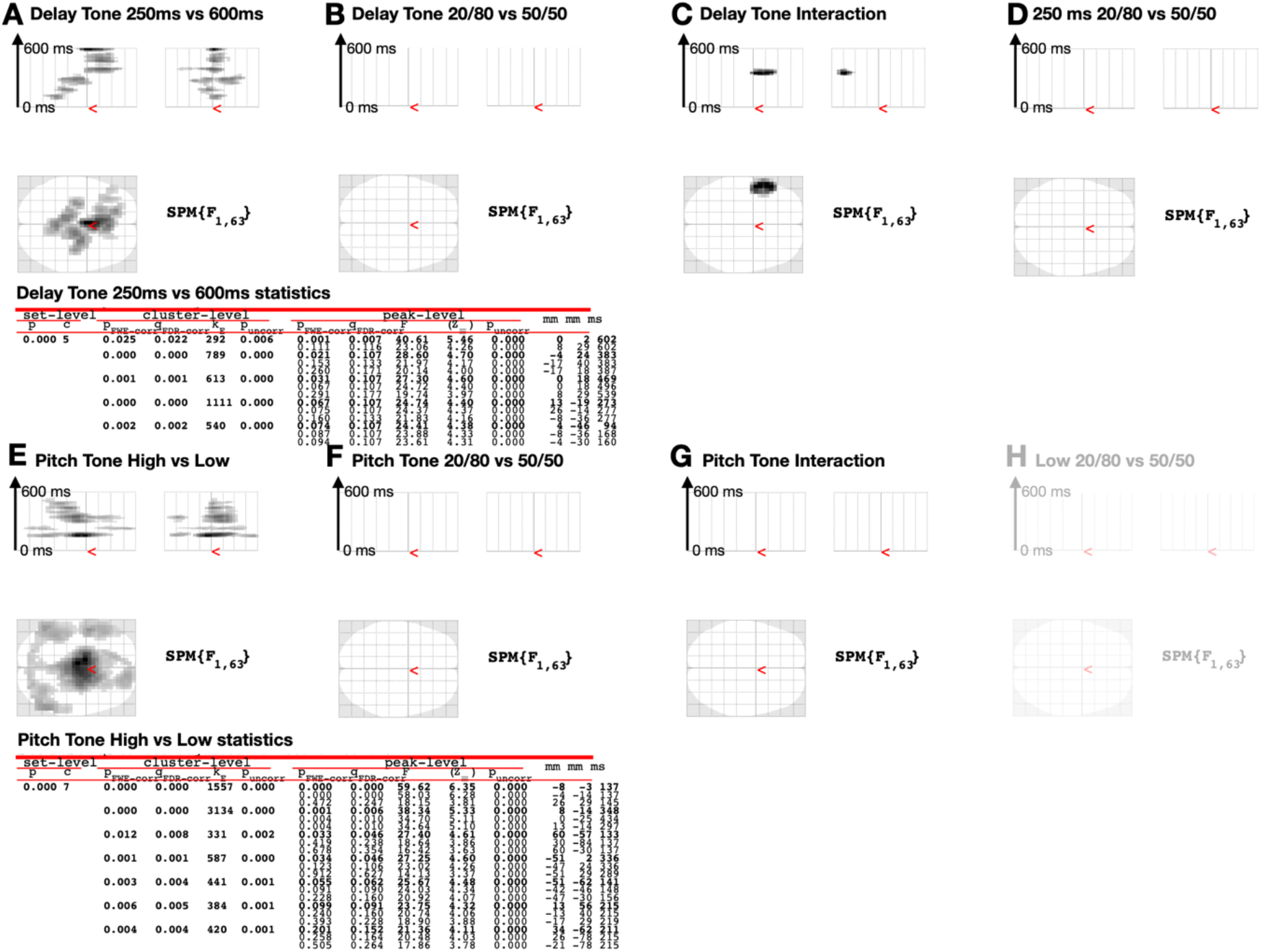
Scalp-time maps of aERPs. **A** Scalp-time map of the main effect of tone delay (250 ms vs. 600 ms) in the Tone Delay experiment, showing significant effects in various medial and frontal regions at different times. **B** No significant time-scalp difference between of the main effect of tone probability (20/80 vs 50/50) in the Delay experiment. **C** Significant interaction between tone delay (250ms vs 600 ms) and probability (20/80 vs 50/50 ms) in a region in the left frontal hemisphere around 350 ms after the tone is played. The interaction between probability and delay reveals differences (p<0.001 uncorrected) in a fronto-medial region 297 ms after the tones are played, and differences in a left lateralised region 172 ms after the tones are played. **D** No significant differences in the scalp-time maps when comparing the identical tones at delay 250m from the 20/80 and 50/50 conditions. **E** Significant main effect of the pitch (High vs Low) for the Tone Pitch experiment in different regions at different times. **F** No significant main effect of tone probability (20/80 vs 50/50) in the Tone Pitch condition. **G** No significant interaction effect between probability and Pitch for the Tone Pitch Condition. SPM{F} map is initially thresholded at p<0.001 uncorrected and then clusters of activated pixels that survive a correction for multiple comparisons p<0.05 FWE are depicted. **H** show the comparison between Low 20/80 vs. 50/50, which due the absent significant interaction in principle is not allowed to be made, and hence for display reasons it is depicted in transparent view.

All scalp time maps for both analyses of MRCPs and aERPs were thresholded at p<0.001 uncorrected for multiple comparisons as cluster forming threshold and only clusters that survive a threshold of p<0.05 corrected for multiple comparisons using gaussian random field theory at the cluster level are displayed.

### Dynamic causal modelling of induced responses

To investigate the effect of the 20/80 and 50/50 conditions on effective connectivity between the inferior parietal cortex (IPC) and preSMA, a DCM analyses of induced responses was conducted on the EEG data from 250 ms before the button press until 600 ms after the button press. EEG data from the analyses of the auditory evoked potentials were used for these analyses. Two separate analyses were performed on the Delay experiment and the Pitch Experiment. The data were further high pass filtered at 4 Hz. Based on the epoched data, 9 different DCM for induced responses were constructed where an equivalent current dipole (ECD) is fitted to the spatial locations used in the model. These comprised two cortical sources, one centred at right IPC, MNI-coordinate: [60, -50, 18], and one centred at right preSMA, MNI-coordinate: [12, 36, 56]. To construct the phenomenological cross-frequency coupling models, the sources are transformed into time-frequency responses using a Morlet wavelet transform. The two areas were coupled bidirectionally in all models 9 models, i.e. the A-matrices of the DCMs. The two different conditions could influence either the couplings from IPC to preSMA bidirectionally, or unidirectionally, i.e. either from IPC to preSMA or from preSMA to IPC alone. Finally, input to the models could either be to both sources, or one of the two sources separately. These models are shown in Figure 5A. After the 9 different models were fitted to the data of each individual participant, a Bayesian model selection (BMS) was performed to select the model that fitted the data best. When the winning model was found, Frequency-Frequency maps of the coupling strengths from the B-matrices were constructed for the 20/80 and 50/50 conditions separately. The difference between these maps were then tested in a repeated measure one-way ANOVA.

**Figure 5:**
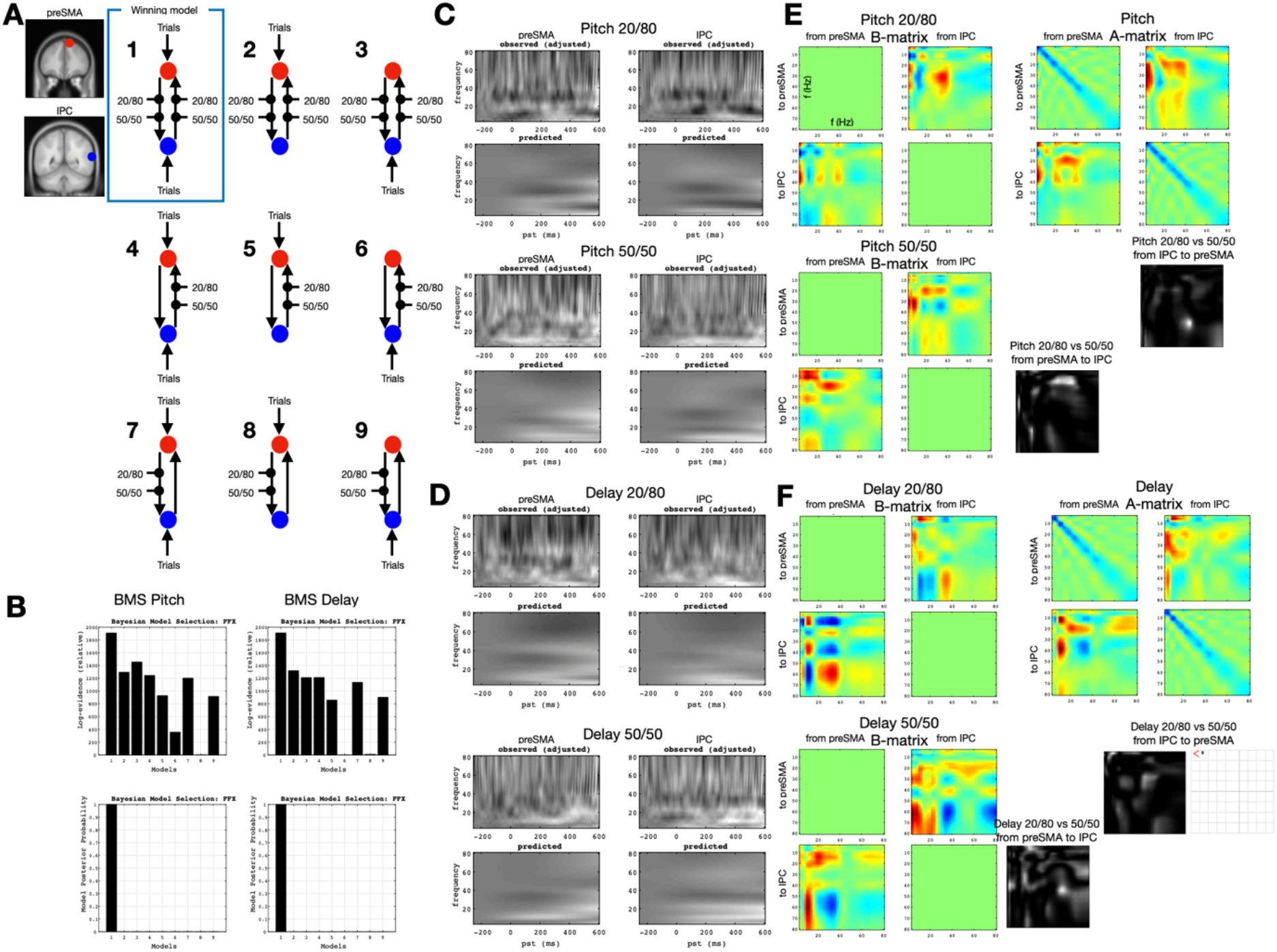
Dynamic causal modelling of induced responses. **A** shows the nine different DCMs that were compared in each of the two experiments, each with sources in preSMA and IPC with locations depicted on a template MRI. **B** shows the results of the Bayesian model selection from the Pitch and Delay experiment, with the Log evidence above and posterior probability below. In both experiments the BMS favour Model 1. **C** and **D** show example of the observed time-frequency decompositions from the two investigated regions preSMA and IPC for the different probability conditions form the Pitch (C) and Delay (D) experiments, respectively. Below is the predicted time-frequency plot based on the outcome of the fitted DCM. **E** and **F** shows the B-matrices, i.e. the task depends on modulation of cross-frequency coupling between the regions of the DCM was well as the A-matrix, which is the task-independent cross-frequency coupling of the DCM. The A and B matrices presented are examples form a single participant’s fitted DCM. In grey-scale images are the F-maps of the cross-frequency couplings comparing the 20/80 and 50/50 conditions for the two separate experiments. For the Delay experiment, the comparison of the coupling from IPL to preSMA in the Delay experiment revealed a single frequency point (4 Hz to 12 Hz), depicted in the grid-display as a single black point, which is uncorrected significant voxel level (p<0.001), but it is not significant at the FWE (p<0.05) corrected level at the cluster level.

## Results

### Behavioural results

The inter press interval (IPI) is calculated for all button presses in the two experiments in each of the probability conditions independently.

We find significant main effects on the IPI of the Condition (Delay: 6.61 s vs. Pitch: 6.33 s, p<2.29×10^−10^, F_1,19_=40.27) and a significant main effect on the IPI of the Probability (20/80: 6.65 s vs. 50/50: 6.29 s, p<8.07×10^−16^, F_1,19_=65.03). We also find a significant interaction of the Condition and Probability on IPI (p<0.0016, F_1,19_=10.0).

Furthermore, doing individual comparisons, we find that Press Delay 20/80 IPI is significantly longer than Press Delay 50/50 IPI (6.72 s vs 6.50 s) and we find that Press Pitch 20/80 IPI is significantly longer than Press Pitch 50/50 IPI (6.58 s vs 6.08 s).

### Press Delay conditions

For the LRP there was no significant difference between neither the early nor the late LRP slope between the 20/80 and 50/50 conditions (See Figure 2A). For the central RP there was also no significant difference between neither the early nor the late central RP slope between the 20/80 and 50/50 conditions (See Figure 2B). Results are displayed in Supplementary Table 1 and 2. The scalp-time maps did not reveal significant (cluster level corrected for multiple comparisons p>0.05 after voxel level threshold set to p<0.001 uncorrected) differences between the Press Delay 20/80 and 50/50 conditions in any scalp region at any time point before the button press (See Figure 2C).

These results show that the probability of the action-effect, when the outcome is manipulated by adjusting the delay of the tone after the button press, is not reflected in differences in neither the early nor late slope of the LRP or RP. The probability of the action outcome also does not influence time-scalp potentials prior to the button press.

### Press Pitch conditions

For the lateralized RP there was no significant difference between neither the early nor the late lateralized RP slope between the 20/80 and 50/50 conditions (Figure 2D). For the central RP there was also no significant difference between neither the early nor the late central RP slope in between the 20/80 and 50/50 conditions (See Figure 2E). See values and statistics in Supplementary Table 3 and 4. The scalp-time maps did not reveal significant (cluster level corrected for multiple comparisons p>0.05 after voxel level threshold set to p<0.001 uncorrected) differences between the Press Delay 20/80 and 50/50 conditions in any scalp region at any time point before the button press (See Figure 2F).

These results show that the probability of the action-effect, when the outcome is manipulated by adjusting the pitch of the tone after the button press, is not reflected in differences in neither the early nor late slope of the LRP or RP. The probability of the action outcome also does not influence time-scalp potentials prior to the button press.

### Delay tone condition

When comparing the latency and amplitude of both the N1, P2, P3 and N4 components of the tones played with 250 ms delay after the button press, we find no significant differences as displayed in Figure 3B and with full statistics in supplementary Table 5. When looking at the scalp-time maps of the main effect of tone delays (Figure 4A) we find several central and lateral areas at varying time intervals that show significant differences between the 250 ms delay tones and 600 ms delay tones. The main effect of probability, i.e. 20/80 vs 50/50 did not reveal any significant differences. There was a significant interaction in a fronto-medial area peaking at 297 ms after the tone was played. However, when comparing only the two identical tones played with 250 ms delay at 80% probability or 50% probability, there is no significant difference in the scalp-time maps.

These results show that there are significant differences in the ERPs when tones are played with two different delays of 250 ms or 600 ms. When we compare two tones played at the same interval (250 ms) with different probabilities either at 50% or 80%, we do not find any significant differences in the ERP.

### Pitch tone condition

When comparing the latency and amplitude of both the N1 and P2 components of the two low pitch tones played with 250 ms delay after the button press under the two different probability conditions, we find no significant differences as displayed in Figure 3D and with full statistics in supplementary Table 6. The scalp-time maps reveal that there is significant main effect of tone pitch, i.e. low vs high pitch tones over central and posterior-lateral regions at multiple times (Figure 4E). These differences can also be seen on the central aERP displayed in Figure 3C, in particular for the high pitch tone played in the 20/80 probability condition, where it serves as an infrequently played odd-ball sound giving rise to clear deflections of the scalp potential away from the identical high pitch tone played in the 50/50 condition. However, when testing the main effect of probability, and the interaction between probability and pitch, there are no significant effects on the scalp-time maps, as displayed in Figure 4F and 4G. Figure 4H display the comparison of the identical low pitch tones played in either the 20/80 or 50/50 conditions, showing no significant difference. This comparison is only for display reasons since no significant interaction was present (i.e. Figure 4G).

These results show that there are significant differences in the ERPs when tones are played with two different pitches. When we compare two tones played with same pitch with different probabilities either at 50% or 80%, we do not find any significant differences in the ERP. We also show that the infrequently played tone does give rise to differences in specific components of the central ERP as well as in multiple potentials at different intervals spread across the scalps, and not only over central regions.

The test of the peak latency and amplitude of the negative MRCP (N-10) around the time point was also compared between the two probability conditions in the Press Delay conditions and also revealed no significant differences neither for the RP nor the LRP as shown in Supplementary Table 7.

The scalp-time maps revealed no significant differences (F-test, p<0.001 uncorrected, cluster p<0.05 uncorrected) between the Press Pitch 20/80 and 50/50 conditions.

These two latter results show that changes in probability of the outcome tone in the pitch experiment are not reflected in differences in the MRCP or LRPs.

### DCM of induced responses

Based on the 9 different models which were identical for both the Pitch and Delay experiment (See Figure 5A), the BMS was employed and for both experiments favoured Model 1 (see Figure 5B). Model 1 is the model with inputs to both sources and where the conditions can modulate both the connection from IPC to preSMA and the connection from preSMA to IPC. The DCMs were based on the time-frequency decompositions (see Figure 5C&D, which show examples from one participant). From the favoured DCM, frequency-frequency maps of the coupling between and within the two regions (A-matrices in Figure 5E&F), were derived. Furthermore, task dependent modulations (B-matrices) of the coupling between the two regions were also derived from the models as frequency-frequency coupling maps. When these are compared for the coupling in both directions, there are however no significant differences (p<0.001 uncorrected at voxel level and p<0.05 family wise error corrected at cluster level) for any of the comparisons between the 20/80 and 50/50 conditions in both experiments for the task dependent couplings. The comparisons are displayed as raw F-maps of the frequency-frequency coupling maps. The comparison of the coupling from IPL to preSMA in the Delay experiment revealed a single frequency point (4 Hz to 12 Hz) p<0.001 uncorrected significant voxel, but it is not significant at the FWE cluster corrected level (p<0.05).

These DCM results show that the coupling between IPC and preSMA is not sensitive to changes and probability of the outcome of an action, and thereby not a possible electrophysiological marker of binding between action and outcome.

## Discussion

The overall finding of this study is that we find no evidence of EEG correlates related to a temporal binding of actions and effects. Despite a previous study that has found electrophysiological indications of measures that reflect the intentional binding phenomenon (Jo et al. 2014) and related processes such as the positive experience of agency (Timm et al. 2016), it was not possible to show a relationship between a simple action-effect task, where the probability of an action leading to a specific effect was subjected to a modulatory factor to increase or decrease binding. This approach has previously been successful in modulating intentional binding (Engbert & Wohlschläger 2007; Moore and Haggard 2008; Moore et al. 2009; Voss et al. 2010) in behavioural experiments.

### Accessing behavioural differences

The only significant differences we find is a slightly skewed temporal distribution of the inter trial interval. The interval between consecutive button presses is generally 0.33 s slower in the high probability (20/80) condition compared to the low probability (50/50) condition regardless of whether it was the Pitch or Delay experiment. If the effect were mainly due to the larger proportion of long delays (600 ms) in the 50/50 Delay condition, one would expect to find that the high probability conditions displayed shorter inter trial interval, not longer, and the effect should only be present in the Delay Experiment, not the Pitch Experiment. Another possibility is that the infrequent high pitch tones in the 20/80 pitch condition introduced an effect similar to post-error slowing, i.e. the surprising tones introduced a slight delay. However, as Figure 1E shows, the two experiments showed the opposite effect.

To maintain the possibility of measuring readiness potentials from all trials, without having interference between two consecutive trials, we made a deliberate choice in the experimental design, which does not allow intervals between two consecutive button presses to be shorter than 3 s. That may distort the distribution of inter press intervals, but despite this, the distribution of IPIs follows previous findings (Schurger et al. 2012) using similar procedures.

Interestingly, different actions can be influenced by the outcome condition of this simple act, as shown by Horwáth et al. (2018). Horwáth et al. (2018) show that the interval between two consecutive actions which were followed by a tone, were shorter for pinch and button actions when these were followed by a tone, where as a tap action followed by a tone was performed with longer interval compared to actions alone. Similarly, exerted force was generally higher in conditions that did not have a tone after the action. Unfortunately, we did not record how much force was exerted by the participants when they pressed the button. We however could observe significant differences in the time intervals between each button press, so with a predictable (80%) outcome the IPI was longer than with a less predictable (50%) outcome. However, further investigations would be needed to clarify whether changes in outcome probability would change the actual action performed.

### Accessing readiness potentials

Contrary to Jo et al.’s (2014) findings, there were no differences in the early slope of the RP, neither the central RP nor the LRP between the situation when the outcome of the action of pressing a button was highly predictable (80%) and when it was unpredictable (50%) for both the experiments with different pitches as well as delays of the tones. In addition, we explored differences between the two conditions in terms of the slope of the late RP, the peak amplitude and latency of the MRCPs, here denoted latency and amplitude of the N-10. Here, we did not find any differences between the two conditions for both experiments (Pitch and Delay). To further explore the differences between the higher (80%) and lower (50%) probability conditions, the scalp-time potential maps were tested. These explore any differences in scalp potentials at any location of the scalp between at the time-interval from 1.5 s before the button press until button press. Because of the many areas and time point tested, precautions were made to ensure that multiple comparisons are adjusted appropriately. With such adjustment we find no differences in the potential at any scalp location at any time prior to the button press for any of the two experiments.

The differences found by Jo et al. (2014) where present in an experiment where the judgment of temporal intervals was assessed explicitly using a Libet clock experimental procedure (Libet et al. 1983). Our study was not conducted while participants had to judge either the time of the button press or time of the tone, which in combinations can provide insights into the binding phenomenon. Jo et al. (2014) show that the early slope of the RP is correlated with binding, whereas in the present study we only can test whether there is a difference in slope between the two probability conditions. But if we accept that the intentional binding phenomenon is a general mechanism that is present in all situations where an action causes an effect, we should expect that the underlying neurobiological mechanism was present regardless of whether the binding phenomenon was explicitly assessed. However, given the absence of any difference in LR slope, it may be worth considering whether that finding may be directly related to the task of judging the timing of either the action or the effect.

There is no difference in the amplitude or the delay of auditory tones played in a high probability condition 20/80 compared with a low probability (50/50) condition. Also, there are no differences in neither the early nor late slope of the lateralized (LRP) or central RP between high and low probability conditions.

### Accessing auditory evoked potentials

When investigating the aERPs, we did not find any differences in the amplitudes or latencies of neither the N1 nor P2 components of the aERPs from the two similar tones presented in the high (80%) probability compared to the low (50%) probability conditions. We were therefore not able to replicate the findings of Timm et al. (2016), which related the P2 component to sense of agency. But the method of inducing a sense of agency in the study by Timm et al. (2016) was to adapt an action to an effect by manipulating the interval between the action and the effect, and not the probability of the outcome, as was done in this present experiment. The probability of an action leading to a specific effect has been shown to be important for subjective sense of agency (Moore and Haggard 2008; Moore et al. 2009). We did find significant main effects of the Delay tone, so when comparing the 250 ms Tones with the 600 ms Tones across the two probability conditions for multiple components of the ERP, as revealed by the scalp-time potential comparisons. Klaffhen et al. (2019) investigated differences in both the N1 and P2 component of self-generated and externally generated tones, but this relation was also manipulated in terms of delay after a key press (0 or 750 ms). Whether the differences in using probabilities as opposed to delays may induce differences in the experience of sense of agency and subsequent effects on the various ERP components remain a topic for further investigations. Therefore, the results may not be directly comparable. This lack of comparability however may suggest that different underlying principles are responsible for monitoring temporal delays and probabilities of outcomes. Farrer et al. (2008) have previously shown that delays and spatial distortions in feedback manipulation studies of drawing movements give rise to different subjective reports.

However, the manipulation in the high probability condition does seem to work, since the infrequently (20%) played tone, in particular for the high pitch tone does give rise to large central positive ERPs with delays of approximately 300-400 ms (Schafer & Markus 1973; Duncan-Johnson & Donchin 1977). For the experiment with modulations of the Delay, we do not see the large central aERPs for the infrequently (20%) played tones in the high probability condition. This is mainly because the infrequently played tone is played with 600 ms delay, and hence does not generate the same surprisal response compared to a situation where the frequently played tone had been played with the long 600 ms and the infrequently tone played with 250 ms. When 250 ms has elapsed since the button press and no tone has been heard, it will come after another 350 ms. The reason for including the frequent/infrequent tone with the chosen delays was to be able to compare the frequently played tones also with the tones from the pitch experiment, if it turned out that any of the experiments and conditions revealed interesting results.

Delays between actions and subsequent effects have been used by Imaizumi & Tanno (2019) and Wen et al. (2015) to study intentional binding and sense of agency but with very different results. Imaizumi & Tanno (2019) showed that longer delays between an action and an event decrease explicit judgments of agency and reduce intentional binding. However, Wen et al. (2015) showed the same effect on explicit agency but increased binding effect with longer delays. This suggests that delays cannot consistently account for (a lack of) intentional binding given the difference in results. However, in the present study, the delay was not used to explicitly change the intentional binding effect, but to create high or low probability action-effect contingencies and subsequently focus on the comparison between tones played with the same 250 ms delay.

A number of recent studies (Kleimaker et al. 2020; Takacs et al. 2020; Takacs et al. 2021) have investigated stimulus-response binding with EEG in light of the popular Theory of Event Coding (Hommel et al. 2001). One of the main findings of these experiments is that a cluster of activity patterns reflect the intermediate processes linking the stimulus. This cluster of activity can be observed in relation to the P3 ERP component or a stimulus that precedes the subsequent action. It is questionable whether the absence of any differences between the P3 ERP components, when the outcome of the action is played with two different probabilities, can be interpreted as lack of evidence of the event coding. We rather interpret that our lack of difference is due to the overall difference in experimental procedure when comparing the different approaches. In our experiment, the tone follows the action, and in the studies by Kleimaker et al. (2020) & Takacs et al. (2020; 2021), the action is performed in response to the stimulus of which the P3 ERP component is analysed on. It would be very interesting if stimulus-response binding and action-effect binding are reflected in exactly the same underlying electrophysiological mechanisms. However, to our knowledge, the phenomenon of intentional binding has not been studied with reversed order of action and effect, i.e. in a stimulus-response way. Therefore, further studies are needed to clarify whether both stimulus-response binding and action-effect binding can be accounted for by the same P3 ERP component mechanisms.

The comparison of the MRCP and aERPs was to calculate the temporal interval between the peak MRCP (N-10 component) and aERP. For the N-10 component, we did not use the rectified EMG as reference point, but the time of the button press as indicated by the trigger in the EEG file. Although the exact delay of the N-10 component may differ, the electromechanical delay between the EMG onset and actual button press can be assumed to be constant, and negligible in comparison to the overall interval from N-10 to N1 used for this calculation. This comparison also did not reveal any neural signatures of an attraction in time of the two potentials when they were played in the high probability condition compared with the low probability condition in any of the experiments. The N-10 component is interpreted as being related to pyramidal tract motor neuron activity in primary motor cortex (Shibasaki & Hallett 2006). To our knowledge, there has not been any mentioning of changes in latency of the N-10 component in relation to probabilistic variations in action-effect outcomes, and this comparison that is made here can be considered exploratory.

### Accessing Dynamic Causal models

Finally, the DCM analysis did not reveal any differences in coupling between IPL and preSMA when comparing the high probability or low probability conditions in either of the experiments during the period between the button press and the tone. This negative finding may be a little less surprising, given that the previous study (Ritterband-Rosenbaum et al. 2014) relating sense of agency to IPL – preSMA coupling was performed using a feedback manipulation task rather than a simple action-effect task. In the previous study by Ritterband-Rosenbaum et al. (2014), participants watched a cursor moving on a screen controlled by a pen tablet movement. The displayed cursor movement was then distorted to different degrees to manipulate sense of agency. We consider a feedback manipulation task, an ongoing (visuo)motor task, where presence and absence of sense of agency is modulated with manipulations of the sensory feedback provided during the experiment, for a detailed discussion of this distinction between action-effect and feedback manipulation task, see Christensen & Grünbaum (2017, 2018).

In conclusion, there were no neural signatures that reflected an attraction of EEG signals on any of the known electrophysiological measures related to the intentional binding phenomenon or subjective sense of agency in an experiment that has used manipulations of probabilistic relations between actions and outcomes as an indirect way of probing the intentional binding phenomenon.

### Limitations of the study

The main objection against the findings of this study, or rather the lack of findings, is naturally that we used no explicit measure of intentional binding with which we could compare the electrophysiological measures. This may be the main reason why we did not replicate the finding from Jo et al. 2014. But our omission of an explicit measure was deliberate, as the introduction of the explicit measure of intentional binding should not be the principle cause of the intentional binding phenomenon. The idea behind the intentional binding phenomenon is that it is an ever-present phenomenon, which is present when one feels oneself as the agent of an action (i.e. that intentional binding is related to sense of agency), which is the most predominant idea, or intentional binding may be a phenomenon that links cause and effect to each other. Regardless of whether intentional binding reflect sense of agency or causal linking it should be present regardless of whether it is measured or not.

An important part of experimental designs is that they engage participants so that they maintain focus on the task they are asked to perform. In this experiment, the ecological validity and measure of performance is extremely limited. Participants obtained no feedback on their performance, because there was no explicit measure of performance in this task. This may limit the engagement of participants with the task, and potentially affect results such as a lack of a binding effect. However, provided with the assumption that our voluntary actions are accompanied by a sense of agency (Haggard 2017), regardless of whether we pay attention, it is expected that regardless of whether one makes explicit enquiries, the underlying “buzz” or sense of agency should be present. It is therefore valuable to perform experiments that are not governed by the necessity of explicitly making enquiries into the performance of the action, to study the phenomenon of a sense of agency that accompanies actions. If it requires explicit enquiries into the actions performed, it could suggest that a positive sense of agency is only present when one explicitly pays attention to it. In contrast, a negative or lack of sense of agency can be elicited by both limited attention to the task or lack of experience of a cause and effect relationship independent of voluntary involvement.

The use of different probabilistic relations between the action and its outcome is the other critical issue in the design of this experiment. To appreciate the underlying idea of this experiment, we rely on a number of mappings between relationships. One is to accept that changing probabilities of action-effect outcomes causes differences to the intentional binding phenomenon. This has been shown in previous studies that high action-effect outcome probability gives high binding effect, i.e. shorter perceived temporal interval between action and effect (Engbert & Wohlschläger 2007; Moore and Haggard 2008; Moore et al. 2009; Voss et al. 2010). Another important relation is the voluntary/involuntary action and how this relates to intentional binding. Haggard et al. (2002) showed how voluntary actions are associated with binding and involuntary with repelling. This finding, that manipulating with the action-outcome probability does not lead to electrophysiological bindings, may suggest that the binding phenomenon requires active enquiry to be present, and it questions whether it is a phenomenon that is ever-present in our everyday actions.

As presented in the introduction, a number of recent studies (Saito et al. 2015; Schwarz et al. 2019; Kirsch et al. 2019; Ma et al. 2020) have not been able to establish a relation between sense of agency and intentions with the intentional binding effect, and this assumption has been an important part in the design of this experiment, which was conducted before these studies were published. The result of this study may therefore not be surprising in the light of these recent behavioural studies, which have questioned the relation between sense of agency and intentional binding and among other findings shown that binding can occur in the absence of action intentions.

We acknowledge that these recent findings may be sufficient reasons not to consider the argument that all voluntary actions will give rise to an intentional binding phenomenon, as originally proposed that regardless of whether it is measured or not, the feeling of agency and its accompanying phenomenon of intentional binding should always be present. But since this argument was the starting point for this study, we nevertheless feel obliged to display our results even though the argument could be weakened by the latest experimental findings.

### Construct validity

To establish construct validity of the concept of sense of agency in simple action-effect studies with experimental procedures like the intentional binding phenomenon, subjective report and electrophysiological measures, it is necessary that relationships between the various procedures and measures show consistency.

To establish construct validity concerning action-effect studies with sense of agency, intentional binding and electrophysiological measures, Figure 6 provides insight into how this can be established, and what this study adds to that discussion (for extended discussion see Grünbaum & Christensen (2020)). It appears that there are consistent relations between action-effect (A-E) and sense of agency, sense of agency and intentional binding, sense of agency A and ERPs, and A-E and ERP when the manipulation is differences in delays between the action and the event. However, the relationship between intentional binding and electrophysiological measures even with delays as experimental manipulation remain unresolved. Using probabilistic relations between actions and effects does not seem to be a viable solution to establish complete construct validity of all types of manipulations. The study that we present here displays a missing relationship between action effect manipulations and electrophysiological measures.

**Figure 6:**
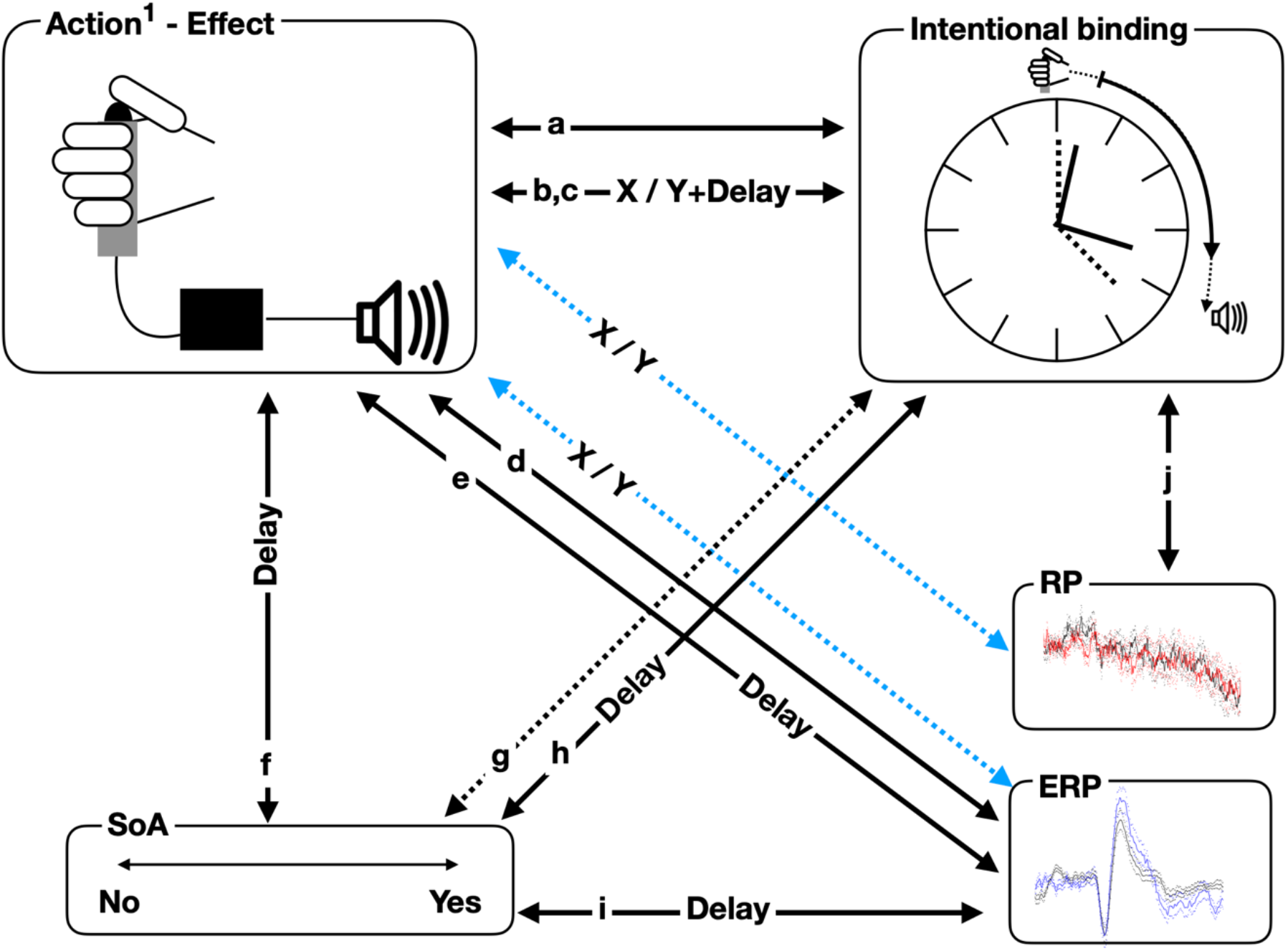
Overview of established and missing relations ships between action-effect tasks, intentional binding, sense of agency and electrophysiological measures. 1) Actions can either be voluntary or involuntary, TMS induced twitches is an example of an involuntary action, for instance as used by Haggard et al. (2002). The black box in the Action-Effect box indicates that various relationships are used as experimental manipulations. Fully drawn arrows between two boxes indicate a relationship between two procedures and measures have been identified. Dotted arrows between two boxes indicate that a relationship has been tested but not identified. If the arrow is named ‘Delay’ it means that the introduction of a variable delay between the action and the effects is the main reason for modulating the relationship. For instance, longer delay leads to decreased sense of agency (SoA). If the arrow is named ‘X/Y’ it means that the introduction of variable probabilities between actions and effects modulate the relationship. Blue dotted arrows indicate this present study. a) Haggard et al. (2002), b) Moore & Haggard (2008), c) Engbert & Wohlschläger et al. (2007), d) Schafer & Markus (1973), e) Timm et al. (2016), f) Sato & Yasuda (2005), g) Saito et al. 2015 + Schwarz et al. 2019 + Kirsch et al. 2019 + Ma et al. 2020, h) Imaizumi & Tanno (2019), i) Jo et al. (2014).

## Conclusion

It was not possible to identify any electrophysiological measures that are related to higher probability action-effect relationships compared with lower probability relationships, and hence it was not possible to identify electrophysiological markers of intentional binding. Changing the probabilistic relations between an action and its outcome does not seem to be a viable solution to create intentional binding signatures, i.e. a temporal attraction of action and event, but it is effective to create strong neural signatures of irregular relations, similar to odd-ball paradigms. To establish a coherent relationship between action-effect experiments, intentional binding, sense of agency and electrophysiological measures, using probabilistic relations are inferior to the introduction of delays between an action and its effect. Whether delays and probabilistic relations between actions and effects tap into the same underlying neurophysiological mechanism remain so far unknown and requires further investigation.

## Acknowledgements

The study was funded by The Danish Council for Independent Research – Humanities (12-126343). Max Seignette was supported by an Erasmus EU exchange program. MSC was also funded by the Elsass Foundation.

## Funding influences

The funding bodies had no influence on decisions made with respect to where to publish the manuscript. The funding bodies had no influence on any scientific decisions made in the studies presented in this manuscript.

## Author contributions

Idea and design: MSC, Data acquisition: MS, MSC, Data analysis: MSC, MS, Manuscript writing: MSC. All authors approved the final version of the manuscript. We would like to thank Mads Jensen for comments to a preliminary version of the manuscript and useful suggestions for improvements.

---

**Table 1:** Slopes of the early and late components of the LRP and RP for the 20/80 and 50/50

|  | <u>Slope</u> | <u>SD</u> | <u>t-statistics</u> | <u>p-value</u> |
| --- | --- | --- | --- | --- |
| ***Press Delay LRP*** |  |  |  |  |
| *Early 20/80* | -0.0648 $\mu\text{V/s}$ | 0.1866 $\mu\text{V/s}$ | $t_{19} = 0.2042$ | $p = 0.8404$ |
| *Early 50/50* | -0.0838 $\mu\text{V/s}$ | 0.3659 $\mu\text{V/s}$ | | |
| *Late 20/80* | -0.6739 $\mu\text{V/s}$ | 0.6102 $\mu\text{V/s}$ | $t_{19} = -0.1616$ | $p = 0.8733$ |
| *Late 50/50* | -0.6479 $\mu\text{V/s}$ | 0.6563 $\mu\text{V/s}$ | | |
| ***Press Delay RP*** |  |  |  |  |
| *Early 20/80* | -0.0476 $\mu\text{V/s}$ | 0.1760 $\mu\text{V/s}$ | $t_{19} = 1.0271$ | $p = 0.3173$ |
| *Early 50/50* | -0.1521 $\mu\text{V/s}$ | 0.3969 $\mu\text{V/s}$ | | |
| *Late 20/80* | -0.3946 $\mu\text{V/s}$ | 0.7875 $\mu\text{V/s}$ | $t_{19} = -0.3453$ | $p = 0.7336$ |
| *Late 50/50* | -0.3391 $\mu\text{V/s}$ | 0.6846 $\mu\text{V/s}$ | | |

**Table 2.** Amplitude and latency of the late components of the LRP and RP around the time of button press in the Press Delay task for the 20/80 and 50/50 conditions.

|  | <u>Latency</u> | <u>SD</u> | <u>t-statistics</u> | <u>p-value</u> |
| --- | --- | --- | --- | --- |
| ***Press Delay LRP*** |  |  |  |  |
| *20/80* | -8.4 ms | 58.2 ms | $t_{19} = 0.3740$ | $p = 0.7126$ |
| *50/50* | -15.2 ms | 63.0 ms |  |  |
| ***Press Delay RP*** |  |  |  |  |
| 20/80 | 10.2 ms | 51.6 ms | t<sub>19</sub>= 1.7039 | p=0.1047 |
| 50/50 | -18.9 ms | 60.5 ms |  |  |
|  | <u>Amplitude</u> | <u>SD</u> | <u>t-statistics</u> | <u>p-value</u> |
| ***Press Delay LRP*** |  |  |  |  |
| 20/80 | -0.7791 μV | 0.6365 μV | t<sub>19</sub>=0.3485 | p=0.7313 |
| 50/50 | -0.8562 μV | 0.7958 μV |  |  |
| ***Press Delay RP*** |  |  |  |  |
| 20/80 | -0.6841 μV | 0.6709 μV | t<sub>19</sub>=0.2594 | p=0.7981 |
| 50/50 | -0.7338 μV | 0.7710 μV |  |  |

**Table 3:**
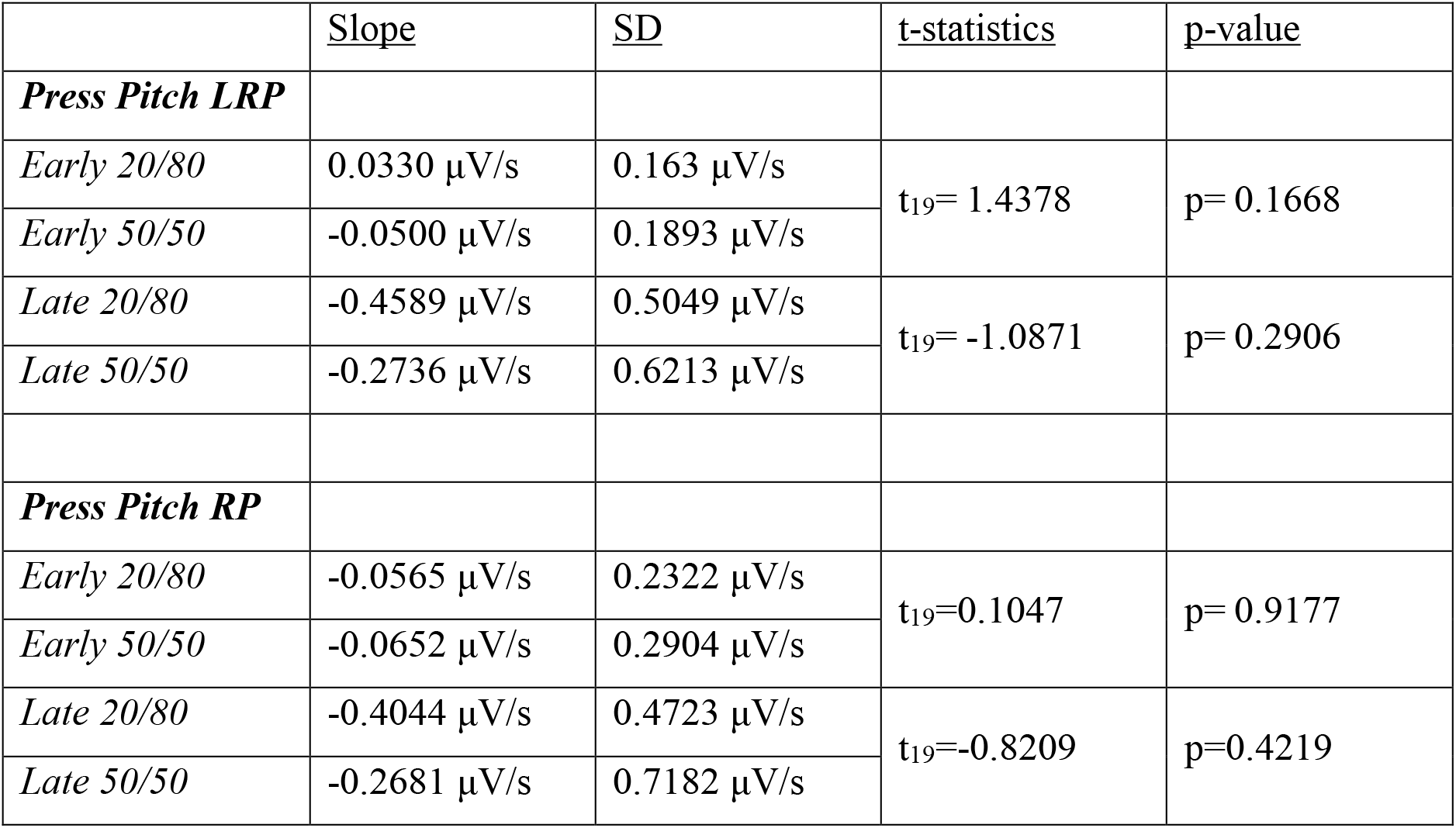
Slopes of the early and late components of the LRP and RP for the 20/80 and 50/50 conditions of the Press Pitch task.

**Table 4.** Amplitude and latency of the late components of the LRP and RP around the time of button press in the Press Pitch task for the 20/80 and 50/50 conditions.

|  |  |  |  |  |
| --- | --- | --- | --- | --- |
|  | <u>Latency</u> | <u>SD</u> | <u>t-statistics</u> | <u>p-value</u> |
| **Press Pitch**<br>**LRP** |  |  |  |  |
| 20/80 | -3.5 ms | 67.3 ms | $t_{19} = -0.3284$ | $p = 0.7462$ |
| 50/50 | 2.0 ms | 64.0 ms |  |  |
| **Press Pitch RP** |  |  |  |  |
| 20/80 | -1.2 ms | 54.4 ms | $t_{19} = -0.8981$ | $p = 0.3804$ |
| 50/50 | 13.7 ms | 53.8 ms |  |  |
|  | <u>Amplitude</u> | <u>SD</u> | <u>t-statistics</u> | <u>p-value</u> |
| **Press Pitch**<br>**LRP** |  |  |  |  |
| 20/80 | -0.6581 $\mu V$ | 0.5999 $\mu V$ | $t_{19} = -0.2109$ | $p = 0.8352$ |
| 50/50 | -0.6172 $\mu V$ | 0.5814 $\mu V$ | | |
| **Press Pitch RP** |  |  |  |  |
| 20/80 | -0.7106 $\mu V$ | 0.5489 $\mu V$ | $t_{19} = 0.1815$ | $p = 0.8579$ |
| 50/50 | -0.7409 $\mu V$ | 0.4920 $\mu V$ | | |

**Table 5:** Latency and amplitude of the N1, P2, P3 and N4 components of the aERP in the Delay experiments.

|  |  |  |  |  |
| --- | --- | --- | --- | --- |
|  | <u>Latency</u> | <u>SD</u> | t-statistics | p-value |
| **250 ms delay tone** |  |  |  |  |
| *N1 20/80* | 110.2 ms | 8.4 ms | $t_{19} = -0.227$ | $p = 0.823$ |
| *N1 50/50* | 110.5 ms | 9.1 ms |  |  |
| *P2 20/80* | 187.9 ms | 24.0 ms | $t_{19} = 0.657$ | $p = 0.519$ |
| *P2 50/50* | 185.0 ms | 17.4 ms |  |  |
| *P3 20/80* | 293.2 ms | 35.9 ms | $t_{19} = 0.1396$ | $p = 0.890$ |
| *P3 50/50* | 292.8 ms | 28.6 ms |  |  |
| *N4 20/80* | 436.7 ms | 34.5 ms | $t_{19} = 0.0$ | $p = 1.000$ |
| *N4 50/50* | 436.7 ms | 45.3 ms |  |  |
|  | <u>Amplitude</u> | <u>SD</u> | t-statistics | p-value |
| **250 ms delay tone** |  |  |  |  |
| *N1 20/80* | -1.2673 μV | 0.7246 μV | $t_{19}=1.961$ | $p=0.065$ |
| *N1 50/50* | -1.7531 μV | 0.9307 μV | | |
| *P2 20/80* | 1.6255 μV | 0.9433 μV | $t_{19}=-1.611$ | $p=0.124$ |
| *P2 50/50* | 1.915 μV | 0.9937 μV | | |
| *P3 20/80* | 0.6704 μV | 0.6954 μV | $t_{19}=0.4223$ | $p=0.678$ |
| *P3 50/50* | 0.6138 μV | 0.5634 μV | | |
| *N4 20/80* | -0.5843 μV | 0.5209 μV | $t_{19}=2.515$ | $p=0.021$ |
| *N4 50/50* | -0.9940 μV | 0.7143 μV | | |

**Table 6:**
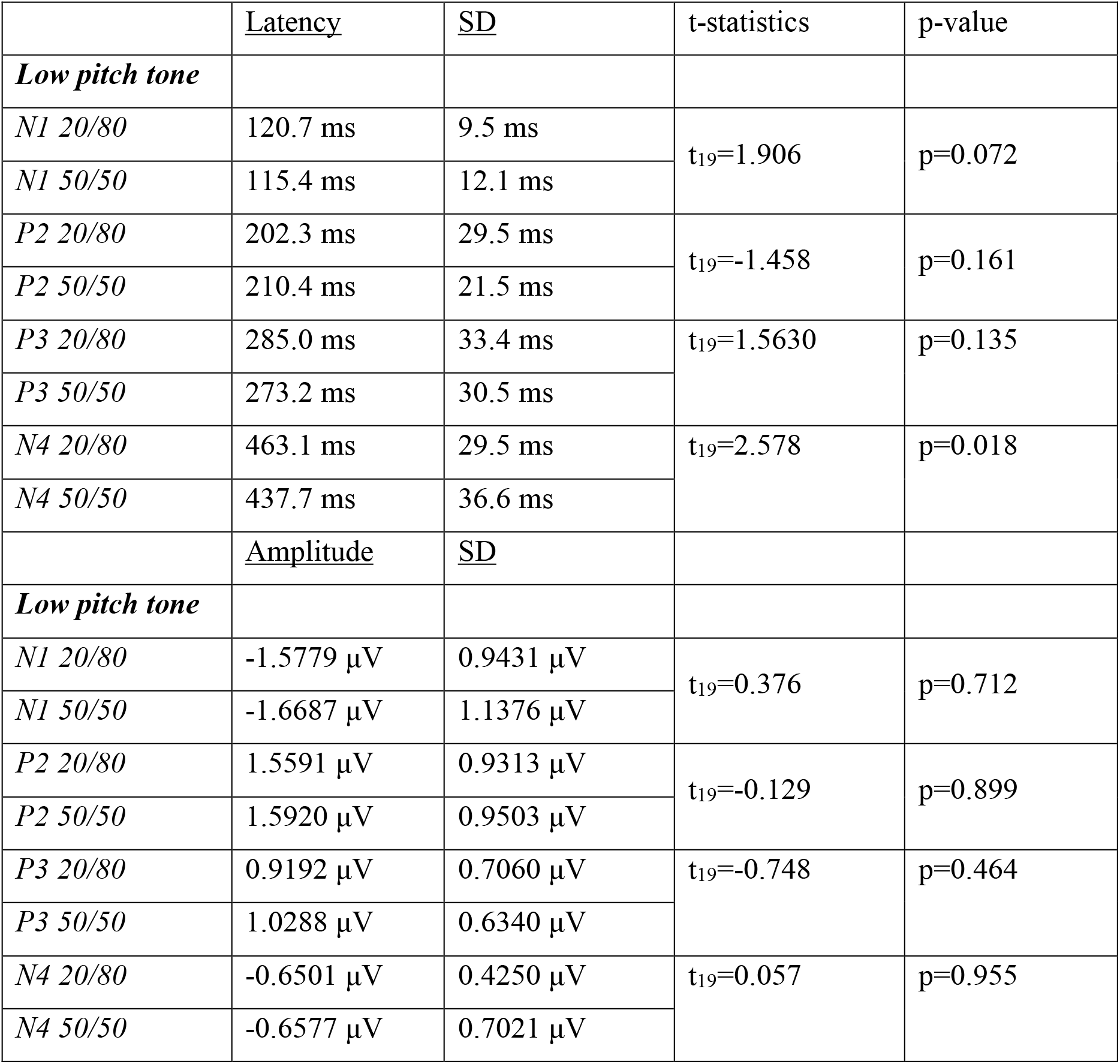
Latency and amplitude of the N1, P2, P3 and N4 components of the aERP in the Pitch experiments.

|  | <u>Latency</u> | <u>SD</u> | t-statistics | p-value |
| --- | --- | --- | --- | --- |
| ***Low pitch tone*** |  |  |  |  |
| *N1 20/80* | 120.7 ms | 9.5 ms | $t_{19}=1.906$ | $p=0.072$ |
| *N1 50/50* | 115.4 ms | 12.1 ms |  |  |
| *P2 20/80* | 202.3 ms | 29.5 ms | $t_{19}=-1.458$ | $p=0.161$ |
| *P2 50/50* | 210.4 ms | 21.5 ms |  |  |
| *P3 20/80* | 285.0 ms | 33.4 ms | $t_{19}=1.5630$ | $p=0.135$ |
| *P3 50/50* | 273.2 ms | 30.5 ms |  |  |
| *N4 20/80* | 463.1 ms | 29.5 ms | $t_{19}=2.578$ | $p=0.018$ |
| *N4 50/50* | 437.7 ms | 36.6 ms |  |  |
|  | <u>Amplitude</u> | <u>SD</u> |  |  |
| ***Low pitch tone*** |  |  |  |  |
| *N1 20/80* | -1.5779 μV | 0.9431 μV | $t_{19}=0.376$ | $p=0.712$ |
| *N1 50/50* | -1.6687 μV | 1.1376 μV | | |
| *P2 20/80* | 1.5591 μV | 0.9313 μV | $t_{19}=-0.129$ | $p=0.899$ |
| *P2 50/50* | 1.5920 μV | 0.9503 μV | | |
| *P3 20/80* | 0.9192 μV | 0.7060 μV | $t_{19}=-0.748$ | $p=0.464$ |
| *P3 50/50* | 1.0288 μV | 0.6340 μV | | |
| *N4 20/80* | -0.6501 μV | 0.4250 μV | $t_{19}=0.057$ | $p=0.955$ |
| *N4 50/50* | -0.6577 μV | 0.7021 μV | | |

## Electrophysiological binding

**Table 7:**
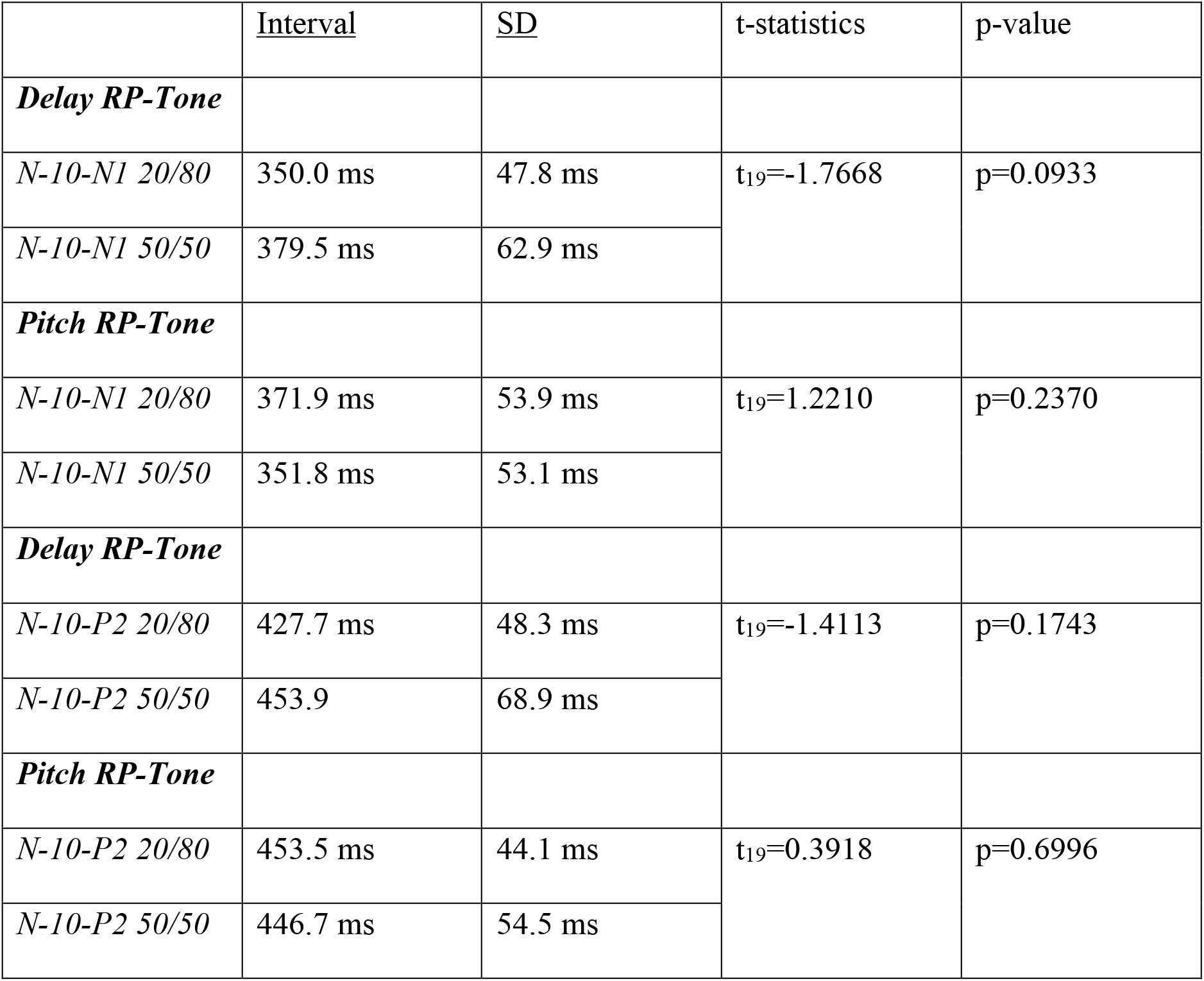
Test of electrophysiological binding, which is the interval in time between the N-10 and respective N1 or P2 ERP component.

|  | <u>Interval</u> | <u>SD</u> | t-statistics | p-value |
| --- | --- | --- | --- | --- |
| ***Delay RP-Tone*** |  |  |  |  |
| *N-10-N1 20/80* | 350.0 ms | 47.8 ms | $t_{19}=-1.7668$ | p=0.0933 |
| *N-10-N1 50/50* | 379.5 ms | 62.9 ms |  |  |
| ***Pitch RP-Tone*** |  |  |  |  |
| *N-10-N1 20/80* | 371.9 ms | 53.9 ms | $t_{19}=1.2210$ | p=0.2370 |
| *N-10-N1 50/50* | 351.8 ms | 53.1 ms |  |  |
| ***Delay RP-Tone*** |  |  |  |  |
| *N-10-P2 20/80* | 427.7 ms | 48.3 ms | $t_{19}=-1.4113$ | p=0.1743 |
| *N-10-P2 50/50* | 453.9 | 68.9 ms |  |  |
| ***Pitch RP-Tone*** |  |  |  |  |
| *N-10-P2 20/80* | 453.5 ms | 44.1 ms | $t_{19}=0.3918$ | p=0.6996 |
| *N-10-P2 50/50* | 446.7 ms | 54.5 ms |  |  |

